# Antiviral Response by Plasmacytoid Dendritic Cells via Interferogenic Synapse with Infected Cells

**DOI:** 10.1101/374496

**Authors:** Sonia Assil, Séverin Coléon, Elodie Décembre, Lee Sherry, Omran Allatif, Brian Webster, Marlene Dreux

## Abstract

Type I interferon (IFN-I) is critical for protection against viral infections. Plasmacytoid dendritic cells (pDCs) massively produce IFN-I against viruses. Physical contacts are required for pDC-mediated sensing of cells infected by genetically distant viruses. How and why these contacts are established remains enigmatic. Using dengue, hepatitis C, zika viruses, we demonstrate that the pDC/infected cell interface is a specialized platform for viral immunostimulatory-RNA transfer, which we named interferogenic synapse and required for pDC-mediated antiviral response. This synapse is an exquisitely differentiated territory with polarized adhesion complexes and regulators of actin network and endocytosis. Toll-like receptor 7-induced signaling in pDCs promotes the interferogenic synapse establishment, thus providing a feed-forward regulation that sustains contacts with infected cells. We propose that the interferogenic synapse is crucial to pDC function as it allows scanning of infected cells to locally secrete IFN-I at the infection site, thereby confining a response potentially deleterious to the host.

**Highlights:**

- pDCs adhere to infected cells via α_L_β_2_ integrin/ICAM-1
- Regulators of actin network and endocytosis polarize at contact
- TLR7-induced signaling potentiates pDC polarity
- Infected cells activate pDCs by interferogenic synapse

## Introduction

Type I interferon (*i.e*., IFNα/β) response is pivotal for protection against viral infections. It is initiated by the recognition of pathogen-associated molecular patterns (PAMPs), including viral nucleic acid, by cellular pathogen recognition receptors (PRRs) such as toll-like receptors (TLRs). This leads to the secretion of type I IFNs along with type III IFNs and inflammatory cytokines, and to the expression of an array of IFN-stimulated genes (ISGs) (Hoffmann et al., 2015). This first line of host response suppresses viral spread and jump-starts the adaptive immune response.

All cells possess signaling pathways designed to trigger antiviral responses against invading viruses. Nonetheless, virtually all viruses have evolved mechanisms to inhibit these host-sensing pathways within cells that they infect (Garcia-Sastre, 2017). For example, dengue virus (DENV) and hepatitis C virus (HCV) encode non-structural viral proteases, which inactivate the adaptors of antiviral pathways induced by different sensors (Aguirre et al., 2017; Aguirre et al., 2012; Horner, 2014; Yu et al., 2012). Despite viral countermeasures, cytokines and ISGs are detected in infected humans, their levels playing pivotal roles in infection clearance and pathogenicity (Martina et al., 2009; Simmons et al., 2007; Snell et al., 2017). This suggests the existence of alternative pathogen-sensing mechanisms. Along this line, we and others uncovered that plasmacytoid dendritic cells (pDCs) produce robust levels of type I IFN in response to the physical contact with infected cells without being productively infected, thus this response is unopposed by viral products (Webster et al., 2016).

pDCs function as sentinels of viral infections and are the major early source of type I IFNs *in vivo* during viral infections (Swiecki and Colonna, 2015). This response is predominantly triggered *via* recognition of viral RNA and DNA species by endolysosome-localized TLR7 and TLR9, respectively. Conversely, type I IFN production by pDCs contributes to several autoimmune or inflammatory diseases, and pDCs are pivotal in cancers (Swiecki and Colonna, 2015; Tomasello et al., 2014). Despite this large spectrum of regulatory functions of the pDCs, the molecular bases controlling their activation are still largely enigmatic.

We previously demonstrated that immunostimulatory viral RNA can be transferred by non-canonical/non-infectious carriers from infected cells to pDCs, leading to IFN production *via* the TLR7 sensor. These PAMP-carriers include viral RNA-containing exosomes and immature viral particles in the context of HCV and DENV, respectively (Decembre et al., 2014; Dreux et al., 2012). The activation of pDCs by exosomal transfer of viral RNA has been also shown for the sensing of genetically distant viruses (Feng et al., 2014; Wieland et al., 2014).

Importantly, while various types of carrier can transfer PAMPs from infected cells to pDCs, the activation of pDCs by virally infected cells requires physical contact (Webster et al., 2016). For example, HCV infected cells secrete exosomes into the culture supernatant at concentrations that are below an excitatory threshold that could be reached in the intercellular space during cell-cell contact (Dreux et al., 2012). In the context of DENV, viral components (i.e. the surface envelope proteins) accumulate at the cell contact between infected cells and pDCs (Decembre et al., 2014). The cell-cell contact is increasingly recognized as a requirement for pDC-mediated antiviral state triggered by genetically distant RNA viruses (*e.g., Flaviviridae, Picornaviridae, Arenaviridae, Retroviridae, Togaviridae*) and in different species (*e.g*., human, mouse, pig) (Bruni et al., 2015; Decembre et al., 2014; Dreux et al., 2012; Feng et al., 2014; Garcia-Nicolas et al., 2016; Lepelley et al., 2011; Takahashi et al., 2010; Webster et al., 2016; Webster et al., 2018; Wieland et al., 2014), nonetheless how the cell-cell contact mediates pDC response is still largely unknown.

Here, we explored the molecular mechanisms underlying the establishment of contacts between pDCs and infected cells, its modulation and impact on the activation of an antiviral state using DENV, HCV and Zika virus as viral models. Our results uncovered that pDCs established cell adhesion with infected cells *via* α_L_β_2_ integrin complex and ICAM-1. We showed that regulators of actin network polarized at contact sites together with components of the endocytosis machinery. These actin-mediated reorganizations define a remodeled territory acting as a platform for the transfer of PAMP-carriers to the endolysosomal TLR7, leading to a pDC IFN response, which prevents viral spread. We further demonstrated that the molecular rearrangements at contact sites were potentiated by the TLR7-mediated response, which thus acts as a positive feed-forward regulation of pDC activation. The molecular specificity endowing the robust IFN response by pDCs is thought to be due to the constitutively high expression levels and enhanced activity of IRF7 (Bao et al., 2016; Honda et al., 2005; Swiecki and Colonna, 2015). Here, we demonstrated that the formation of interferogenic synapse is a pDC attribute critical for their robust IFN response.

## Results

### The α_L_β_2_ integrin/ICAM-1 adhesion molecules are involved in pDC activation by contacts with DENV infected cells

pDC response to viruses needs contact, so we sought to define the molecular bases for establishment of physical contacts between pDCs and infected cells. We tested whether cell adhesion complex can establish pDC/infected cell contact leading to pDC activation. We analyzed the surface expression of a set of representatives of the main cell adhesion/homing families, known to be key for pDC functions, *e.g*., migration and antigen presentation (Sozzani et al., 2010; Swiecki and Colonna, 2015). The L-selectin (also called CD62L), α_L_ integrin (also referred to as ITGAL, CD11a) and intercellular adhesion molecule (ICAM)-1 (also known as CD54) were all expressed at the pDC surface (**Figure 1A and S1A**). ICAM-1 was also present at the surface of the donor cells (*i.e*., DENV infected huh7.5.1 cells as well as uninfected cells), while E-cadherin was present only on the surface of the donor cells (**Figure 1A and S1A**). Next we tested the role of cell adhesion molecules in pDC IFNα production induced by contact with DENV infected cells in antibody-mediated blocking assays (**Figure 1B**). The blockade of L-selectin and E-cadherin had no significant impact on pDC IFNα production induced by DENV infected cells. In sharp contrast, inhibition of both α_L_ integrin and its ligand ICAM-1 (Rothlein et al., 1986; Shimaoka et al., 2003) precluded the pDC IFNα response (**Figure 1B**), in a dose-dependent manner (**Figure S1B**). Viral infectivity and RNA levels were maintained upon similar treatments of infected cells, thus ruling out non-specific effects on DENV replication (**Figure 1B and S1B**). The regulation of the establishment of pDC-infected cell contacts by αL integrin was determined by imaging flow cytometry technology (**Figure S1C-D**). We showed that blocking of α_L_ integrin significantly prevented contacts of pDCs with DENV infected cells (**Figure 1C**).

**Figure 1.**
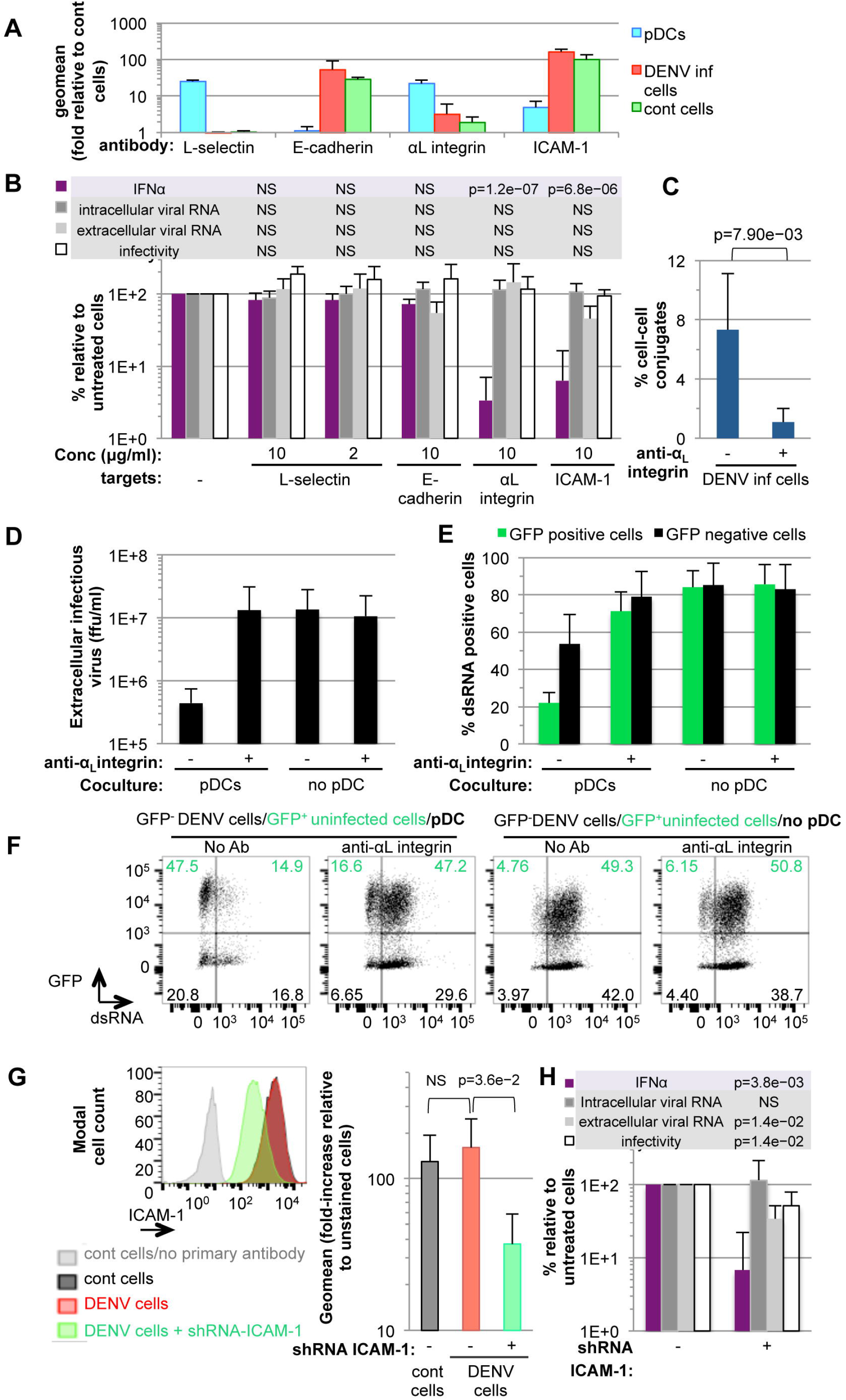
Identification of adhesion molecules involved for pDC-mediated antiviral response against DENV. **(A)** FACS analysis of adhesion molecule expression at the cell surface of pDCs and/or donors cells *i.e*., DENV (inf) infected and uninfected (cont) Huh-7.5.1 cells. Results are presented as fold-increases of surface expression relative to unstained cells or stained only with secondary antibody as reference for immunostained L-selectin or E-cadherin/α_L_integrin /ICAM-1, respectively; 3-4 independent experiments; geomean ± standard deviation (SD); representative histograms in **Figure S1A. (B)** Quantification of IFNα in supernatants of pDCs cocultured with DENV infected Huh-7.5.1 cells that were treated or not with blocking antibodies against L-selectin, E-cadherin, α_L_ integrin or ICAM-1 at the indicated concentrations throughout the course of the coculture. The intracellular and extracellular viral RNA levels and extracellular infectious virus production by DENV infected cells treated with blocking antibodies were analyzed in absence of pDCs but with treatment conditions similar to the analysis of IFN^α^ production in cocultures (*i.e*., incubation time and concentration). Results are expressed as percentages relative to untreated cells, mean ± SD; 4-7 independent experiments. The statistically significances reflected by p-values are indicated in the table; not significant (NS) p-values≥0.05. **(C)** Quantification of conjugates between pDCs and DENV infected cells by Image Stream X technology, as exemplified in **Figure S1C-D**. Celltracer violet stained-pDCs were cocultured with GFP^−^ expressing infected cells for 4-5 hour incubation in presence or not of anti-α_L_ integrin. Results of 5 independent experiments expressed as percentage of conjugates (mean ± SD; n=10 000 events per condition; p-value as indicated). **(D)** Levels of extracellular infectious virus production by DENV infected cells cocultured, or not, with pDCs and treated, or not, with anti-α_L_ integrin (10 μg/mL), for 48 hour incubation. Results are expressed as foci forming unit (ffu) per mL, mean ± SD; 4 independent experiments. **(E-F)** FACS analysis of viral transmission from GFP^−^ DENV infected cells to GFP^+^ uninfected cells cocultured with pDCs (48 hours incubation) and/or in presence of anti-α_L_ integrin (10 μg/mL), as indicated. Results expressed as the frequency of cells positive for dsRNA (a marker of viral replication) in the GFP^+^ and GFP^−^ cell populations, mean ± SD; 4 independent experiments **(E)** and dot plots of one representative experiment (F). **(G-H)** Down-regulation of ICAM-1 expression in Huh7.5.1 cells obtained by transduction using shRNA-expressing lentivirus-based vectors. **(G)** FACS analysis of ICAM-1 surface expression presented with representative histogram and quantifications expressed as fold-increases of geomean relative to unstained cells; n=4 independent experiments; p-value as indicated. **(H)** Quantification of IFNα in supernatants of pDCs cocultured with DENV infected Huh-7.5.1 cells down-regulated or not for ICAM-1 expression, as shown in **G.** The intracellular and extracellular viral RNA levels and infectivity were analyzed. Mean ±SD; n=4 independent experiments.

Next, we demonstrated that the pDC response to contact with DENV infected cells prevented viral propagation. This is shown by the decreased levels of infectious virus production by DENV infected cells when cocultured with pDCs, as compared to the absence of pDC (**Figure 1D**). Of note, the inhibition of α_L_ integrin in infected cell/pDC cocultures fully restored the viral production to levels similar to the absence of pDCs (**Figure 1D**), indicated that the establishment of cell-contact is required for pDC-mediated antiviral response. To further determine the impact on viral transmission, we set up an assay consisting in pDCs cocultured with GFP-positive/uninfected cells and GFP-negative/DENV infected cells. The quantification of the frequency of GFP-positive cells that became infected (*i.e*., positive for dsRNA, a marker of viral replication) at the end of the coculture demonstrated that pDC response readily prevented viral spread from GFP-negative/DENV infected cells to the cocultured GFP-positive (and initially uninfected) cells, and consistently diminished the replication in cells that were infected prior to the coculture *i.e*. GFP positive cells (**Figure 1E-F**). The inhibition of pDC/infected contact via blockage of the α_L_ integrin greatly restored the viral propagation to levels comparable to the absence of pDC (**Figure 1E-F**).

Further, we assessed the impact of the silencing of ICAM-1 expression in infected cells. Reduced ICAM-1 expression (**Figure 1G**) has no impact on virus replication, yet slightly diminishing the release of DENV (**Figure 1H**). In contrast, pDC IFNα production was disproportionally reduced when ICAM-1 expression was silenced in DENV infected cells (**Figure 1H**), further demonstrating the importance of the interaction with ICAM-1 for pDC activation. Together, our results identified α_L_ integrin/ICAM-1 as key cell adhesion complex for the sensing of DENV infected cells by pDC leading to a potent antiviral response.

### Cell polarity of the adhesion molecules at the pDC/DENV infected cells contact

Next we analyzed the subcellular localization of the identified adhesion molecules at the pDC/infected cell interface. We demonstrated that αL integrin specifically accumulated at the pDC/DENV infected cell contacts (*i.e*., over 90% of the analyzed contacts), as opposed to marginal accumulation at contact of CD123 (*i.e*., the α-chain of the interleukin-3 receptor) a surface molecule not involved in cell adhesion (**Figure 2A-C and S2A-B**). The αLβ2 integrin complex (also called lymphocyte function-associated antigen 1; LFA-1) is known to undergo conformational rearrangement to an open form compatible with high-affinity interaction with its ligands, including ICAM-1, and regulated by inside-out signaling (Shimaoka et al., 2002; Shimaoka et al., 2003). The open/high-affinity conformation of α_L_β_2_ integrin complex – detected by monoclonal antibody against the β_2_ I domain epitope (*i.e*., clone m24) (Nordenfelt et al., 2016; Salas et al., 2006; Sen et al., 2018) – polarized at the contact site of pDC/infected cell on the pDC side (**Figure 2D-F and S2C-D**). Of note, cell polarity of α_L_ integrin and the high-affinity conformation of αLβ2 integrin complex was absent when pDCs were not in contact with infected cells (**Figure 2G-H and S2E-F**), thus validating the specificity of the polarization upon cell contact. Further in agreement with the engagement of the α_L_β_2_ integrin complex with its ligand, ICAM-1 (Shimaoka et al., 2002; Shimaoka et al., 2003), the latter was detected at contact on the surface of infected cells apposed to polarized αL integrin on pDC side (**Figure 2I-K**). Our results demonstrate that the open/high affinity conformation α_L_β_2_ integrin complex accumulates at contact facing to its partner ICAM-1.

**Figure 2.**
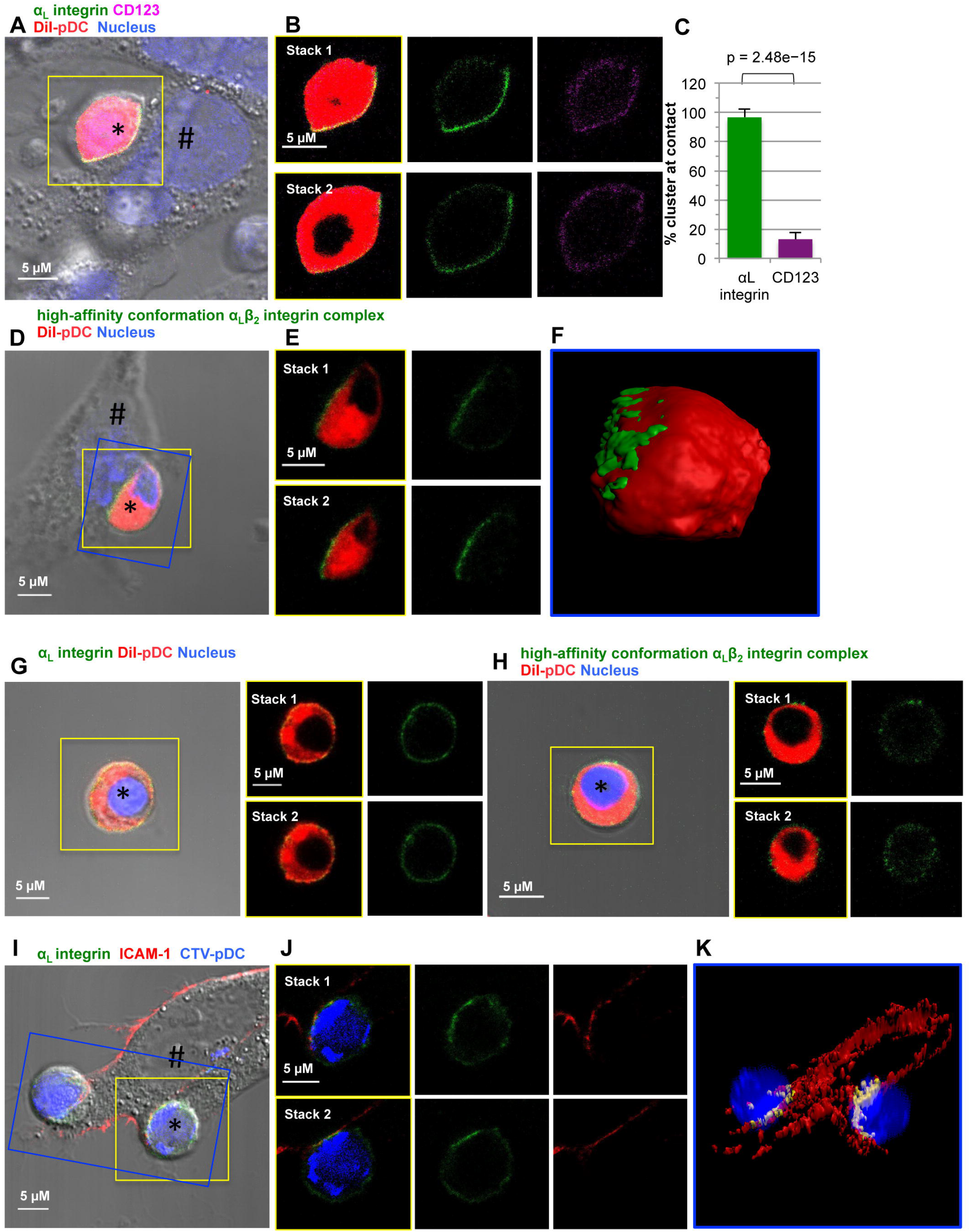
Accumulation of α_L_ integrin and its ligand ICAM-1 at the contact. **(A-C)** Confocal microscopy imaging of immunostained α_L_ integrin (green) and CD123 (purple) in co-cultures of pDCs (stained by the lipophilic dye DiI prior to coculture; red; marked by *) and DENV infected cells (#) for 4-5 hour incubation; nuclei stained with Hoechst (blue). **(A)** Detection of α_L_ integrin, CD123, DiI and Hoechst projected on the phase contrast imaging. **(B)** Magnification of two consecutive Z-stacks of yellow-boxed pDC shown in A. **(C)** Quantification of the frequency of pDCs with accumulation of αL integrin or CD123 at contact. Results of 3 independent experiments are expressed as the percentage of total number of analyzed pDCs in contact with infected cells (mean ± SD; n=39 analyzed contacts; p-value as indicated). **(D-F)** Confocal microscopy imaging of immunostained high-affinity conformation of α_L_β_2_ integrin complex (green) in co-cultures of pDCs (DiI stained; red; *) and DENV infected cells (#), 4-5 hour-incubation; nuclei (Hoechst, blue). Detection of the high-affinity conformation of α_L_β_2_ integrin complex projected on the phase contrast imaging **(D)**, magnification of two consecutive Z-stacks of yellow-boxed pDC shown **(E)** and 3D-reconstruction of consecutive Z-stacks of blue-boxed pDC (F) **(G-H)** Confocal microscopy imaging of immunostained α_L_ integrin **(G)** and the high-affinity conformation of α_L_β_2_ integrin complex **(H)** in pDCs in absence of contact, displayed as in panels **D-E. (I-K)** Confocal microscopy imaging of immunostained α_L_ integrin (green) and ICAM-1 (red) in co-cultures of pDCs (CTV stained; blue; *) and DENV infected cells (#), 4-5 hour-incubation. Staining of α_L_ integrin/ICAM-1 projected on the phase contrast imaging (I) and magnification of two consecutive Z-stacks of yellow-boxed pDC shown (J) and 3D-reconstruction of consecutive Z-stacks of blue-boxed contacts, with colocalized signal in white (K).

### The α_L_β_2_ integrin/ICAM-1 adhesion molecules are required for pDC activation by distinct viruses

The activation of pDCs by HCV infected cells also requires physical contact, yet this involves a distinct PAMP-carrier *i.e*., exosome, while DENV infected cells trigger the pDCs *via* immature viral particles (Decembre et al., 2014; Dreux et al., 2012). We thus tested whether α_L_β_2_ integrin/ICAM-1 adhesion complexes were also required for pDC activation by HCV infected cells. Similar to DENV, blocking of α_L_ integrin significantly prevented contacts of pDCs with HCV infected cells (**Figure 3A**), in keeping with dose-response inhibitions of pDC IFNα production triggered by HCV infected cells (**Figure 3B**). Consistently, blocking of ICAM-1 diminished pDC response against HCV infected cells in dose-response experiments (**Figure 3B**). These results demonstrated that the requirement of α_L_β_2_ integrin/ICAM-1 adhesion molecules for pDC activation is independent of the nature of PAMP-carrier. In sharp contrast, anti-α_L_ integrin and anti-ICAM-1 did not prevent pDC IFNα production triggered by cell-free virus *i.e*., influenza virus (**Figure 3C**), suggesting that α_L_β_2_ integrin/ICAM-1 interaction regulates cell-cell contact mediated-pDC activation.

**Figure 3.**
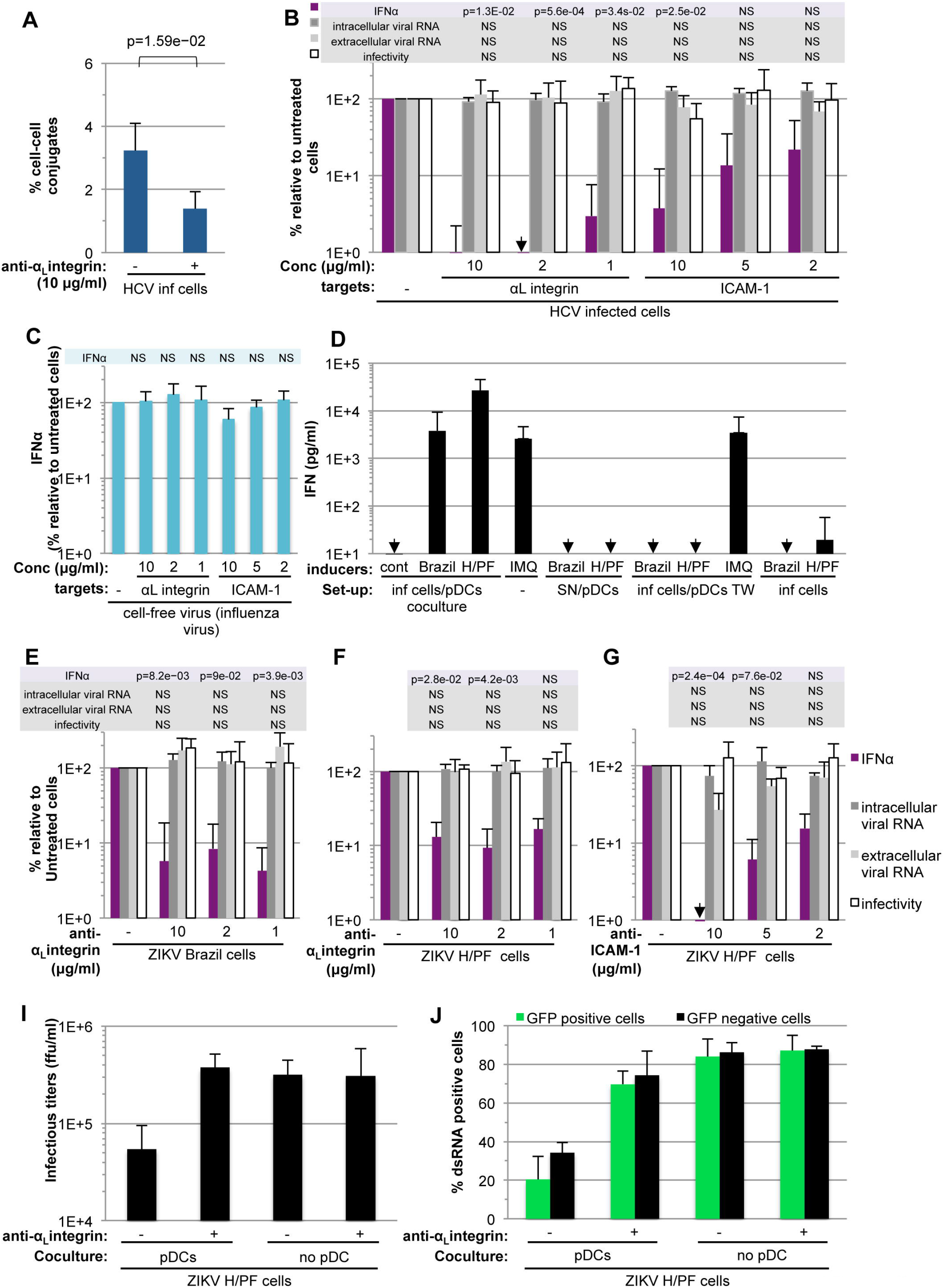
Inhibition of α_L_ integrin and ICAM-1 prevents pDC IFNα production induced by coculture with cells infected by distinct viruses. **(A)** Quantification of conjugate between pDCs and HCV infected cells in presence or not of anti-α_L_ integrin, by Image Stream X technology, as in **Figure 1C, S1C-D**. Results of 5 independent experiments expressed as percentage of conjugates (mean ± SD; n=10 000 events per condition). **(B)** Quantification of IFNα in supernatants of pDCs cocultured with HCV infected cells that were treated or not with blocking antibodies against α_L_ integrin or ICAM-1, at the indicated concentration. The intracellular and extracellular viral RNA levels and infectious virus production by infected cells (in absence of pDCs) treated with blocking antibodies were analyzed. Results and statistical significances are presented as in **Figure 1B**; mean ± SD; n=4-7 independent experiments. **(C)** Quantification of IFNα in supernatants of pDCs stimulated by cell-free influenza virus and treated with blocking antibodies as in **B**; mean ± SD, n=3-4 independent experiments. **(D)** Quantification of IFNα in supernatants of pDCs cocultured with Huh-7.5.1 cells infected by the epidemic ZIKV strains [*i.e*., isolated from patient in Brazil and French Polynesia; H/PF] when cells were in physical contact [coculture], separated by the permeable membrane of transwell [TW], or treated with supernatants from ZIKV infected cells [SN] or imiquimod [IMQ], as positive control. Mean ± SD; n=4-7 experiments; non detectable (arrays). **(E-G)** Quantification of IFNα in supernatants of pDCs cocultured with ZIKV infected Huh-7.5.1 cells treated or not with anti-α_L_ integrin or ICAM-1 blocking antibodies and parallel analysis of intra/extracellular viral RNA levels and infectivity, performed as in panel B. Mean ± SD; n=4-5 **(E)** and n=3 (F-G) independent experiments. **(I)** Quantification of infectious virus production by ZIKV infected cells cocultured with pDCs and treated with anti-α_L_ integrin (10 μg/mL), as in **Figure 1D**. Results are expressed as in **Figure 1D**; 3 independent experiments. **(J)** FACS analysis of viral spread determined as in **Figure 1E-F**. Results expressed as **Figure 1E**; 3 independent experiments.

We thus sought to extend our demonstration to Zika virus (ZIKV), which also represents an important health concern (Miner and Diamond, 2017). Using epidemic ZIKV strains (*i.e*., isolated patients in from Brazil and French Polynesia; H/PF strain), we demonstrated that robust pDC IFNα productions were induced when pDCs were in physical contact with ZIKV infected cells (**Figure 3D**; coculture set-up). On the contrary, no IFNα response was detected when pDCs were exposed to ZIKV infectious supernatants (SN) or upon physical separation of pDCs from ZIKV infected cells by semi-permeable membrane in transwell (TW) experiments (**Figure 3D**). Importantly, we showed that, similar to what is observed for DENV/HCV, pDC IFNα response to ZIKV infected cells is mediated by α_L_ integrin/ICAM-1 adhesion molecules (**Figure 3F-G**).

Next, we demonstrated that, similar to the observation with DENV, pDC response to contact with ZIKV infected cells prevented viral propagation, as determined by the reduced levels of both infectious virus production by ZIKV infected cells cocultured with pDCs as compared to the absence of pDC and viral transmission to GFP-positive cells that were not infected prior to coculture (**Figure 3I-J and S3**). Of note, and similar to DENV, the inhibition of α_L_ integrin in infected cell/pDC cocultures greatly restored both viral production and transmission to levels similar to the absence of pDCs (**Figure 1D**), indicated that the establishment of cell-contact is required for pDC-mediated antiviral response against ZIKV. Together our results demonstrate that the establishment of pDC/infected cell interactions *via* cell adhesion complex is a prerequisite for the antiviral function of pDC in response to cells infected by distinct viruses.

### Actin network and regulators polarization at contacts contribute to pDC response to infected cells

The engagement of integrin family members with their ligand at the cell surface and/or in the extracellular matrix can induce the local recruitment of the actin network (DeMali et al., 2014). This is in line with our preceding observations for the presence of actin at pDC/DENV infected cell contact (Decembre et al., 2014). Here, we showed that actin network accumulated at cell contacts, using both HCV infected cells and the subgenomic replicon (SGR) model for HCV replication (**Figure 4A-C and S4A-C**). HCV SGR cells do not produce virus but trigger pDC IFN response, as previously reported (Dreux et al., 2012; Takahashi et al., 2010) and thus demonstrate that cell polarity at contact is independent of viral production (**Figure 4A-C**). Non-specific polarized immunodetection was ruled out by the simultaneous analysis of another intracellular component, the protein disulfide-isomerase (PDI), an ER marker (**Figure 4A-C and S4A-C**). Additionally, the specificity of actin network polarization at contact was validated by expected decrease of actin clustering upon treatment of pDC/infected cell cocultures with inhibitor of actin polymerization (**Figure 4C**).

**Figure 4.**
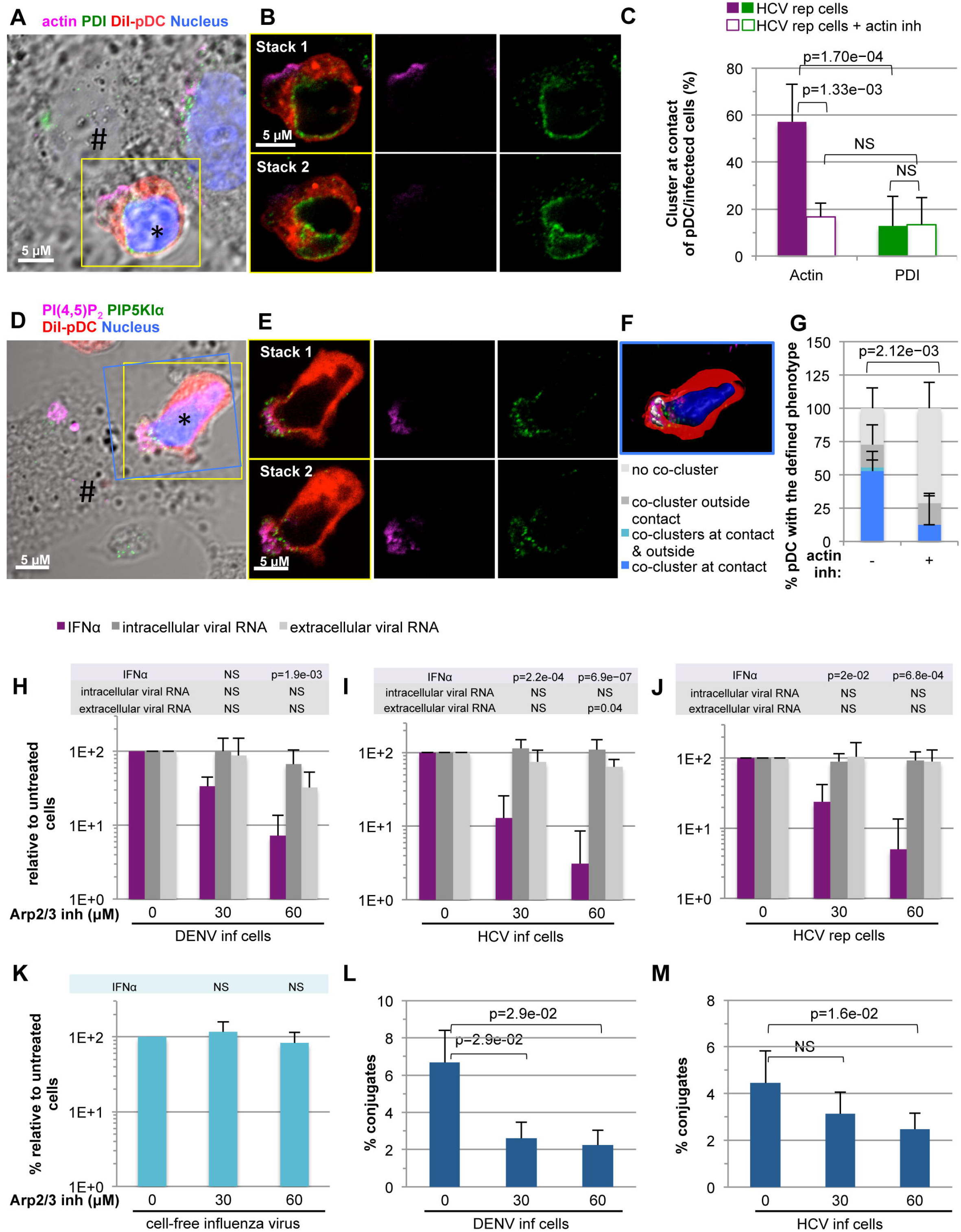
Actin network and regulators polarize at the contact and an intact actin network is required for pDC response. **(A-C)** Confocal microscopy imaging of immunostained actin (purple) and the ER marker PDI (green) in co-cultures of pDCs (DiI stained;*) and HCV SGR replicating cells (HCV rep cells; #), 4-5 hour-incubation; nuclei (Hoechst, blue). Detection of actin and PDI projected on the phase contrast imaging (A), two successive confocal stacks with magnified yellow-boxed pDC (B). **(C)** Quantification of the frequency of pDCs with clustering of actin or PDI at contact in 4-5 hour coculture treated or not with actin polymerization inhibitor (Latrunculin B, 1 mM). Results of 3 independent experiments are expressed as the percentage of analyzed pDCs in contact with HCV rep cells, mean ± SD; n>25 analyzed contacts; p-value as indicated. **(D-G)** Confocal microscopy imaging of immunostained PI(4,5)P_2_ (purple) and PIP5KIα(green) in 4-5 hour cocultures of pDCs (DiI stained, red) and HCV rep cells (#), nuclei (Hoechst). Imaging of PI(4,5)P2 and PIP5KIα projected on the phase contrast **(D)**, confocal stacks of magnification of yellow-boxed pDC **(E)** and 3D-reconstruction of consecutive Z-stacks with colocalization of PI(4,5)P_2_/PIP5KIα in white (F). **G**. Quantification of the phenotype of PI(4,5)P_2_/PIP5KIα coclustering as indicated and including co-cluster at the contacts, outside the contact or no coclustering. Results of 3 independent experiments are expressed as the percentage of total number of analyzed pDCs in contact; mean ± SD; n>21 analyzed contacts; p-value as indicated. **(H-K)** Quantification of IFNα in supernatants of pDCs cocultured with DENV infected cells (H), HCV infected (I), HCV rep cells (J) or stimulated by cell-free influenza virus (K) that were treated or not with inhibitors of Arp 2/3 complex (CK-666, at the indicated concentration). The levels of intracellular and extracellular viral RNA by infected/replicating cells (in absence of pDCs) treated with inhibitors were analyzed. Results and statistical analyses are presented as in **Figure 1B**; mean ± SD; n=4 (H-I) and n=3 (J) independent experiments. **(L-M)** Quantification of pDC/infected cells conjugates by Image Stream X technology. Celltrace Violet stained-pDC cocultured with GFP expressing DENV or HCV-infected cells for 4-5 hours in presence or not of inhibitor of Arp 2/3 complex, as in panels **H-I**. Results of 4 (L) and 5 (M) independent experiments are expressed as percentage of conjugates; mean ± SD; n=10 000 events per condition.

We thus sought to define the subcellular localization of key regulators of actin network scaffolding at the plasma membrane, including phosphatidylinositol 4,5 biphosphate (PI(4,5)P_2_), a plasma membrane-localized modified lipid known to recruit actin-adaptor proteins (Chierico et al., 2014) and type I phosphatidylinositol-4-phosphate-5-kinase α, also called type I phosphatidylinositol phosphate kinase α (referred to as PIP5KIα), which generates PI(4,5)P_2_ from its PI4P precursor (Doughman et al., 2003a; Doughman et al., 2003b). We showed that these regulators co-clustered at the contact (**Figure 4D-E**). The 3D-reconstruction of consecutive Z-stacks further revealed that the co-clusters were present on the side of pDCs (**Figure 4F**). The quantification of PI(4,5)P_2_/PIP5KIα co-clustering revealed that intact actin network was required for the polarization at contact, as demonstrated by the decrease in phenotype when using actin polymerization inhibitor (**Figure 4G**). Of note, PI(4,5)P_2_/PIP5KIα co-clustering was observed at the majority of examined pDC/infected cell contacts, nonetheless the frequency was lower compared to the presence at contact of the αL integrin adhesion molecule (**Figure 2C**). This can reflect the transient nature and/or differential kinetics of these recruitments at contact as compared to the cell polarity of adhesion complexes.

Next, we tested the functional role of actin network regulators, including Arp2/3 complex, required for actin nucleation *via* branching of the actin filaments within network and the Rho GTPase CDC42, which regulates actin polymerization (Rotty et al., 2013). Both are known to be recruited by PI(4,5)P_2_ (Higgs and Pollard, 2000; Rivera et al., 2009; Rozelle et al., 2000). The pharmacological inhibition of either Arp2/3 complex or CDC42 prevented pDC IFNα production induced by either DENV or HCV infected cells (**Figure 4H-J and S4D-E**). This is consistent with similar results obtained upon inhibition of actin polymerization (**Figure S4F**). On the contrary, the levels of intra/extracellular viral RNAs were not, or poorly, reduced in these experimental conditions, thus ruling out unspecific effect on the viral replication of DENV and HCV (**Figure 4H-J and S4D-F**). Of note, similar pharmacological inhibition of Arp2/3 did not prevent pDC IFNα production triggered by cell-free influenza virus (**Figure 4K**), implying that Arp2/3-mediated actin nucleation is pivotal for pDC response induced by contact with infected cells. This assumption was validated by the reduced frequency of pDC/infected cell contacts upon inhibition of Arp2/3 and actin polymerization quantified by imaging flow cytometry (**Figure 4L-M and S4G**), thus inferring that the formation of actin network at contact is pivotal for the establishment and/or reinforcement of the cell contacts. Our results underscore that integrin-mediated adhesion and rearrangement at contact of actin network *via* Arp2/3 and CDC42 regulators, are pivotal for pDC activation triggered by infected cells, presumably by a concerted structuring of the contact site.

### Components of the endocytosis machinery polarize at contacts

We and others previously demonstrated that immunostimulatory viral RNAs is transferred by vesicles (*i.e*., exosome or immature virion) from infected cells to pDCs, thus involving clathrin-mediated endocytosis by pDCs (Decembre et al., 2014; Dreux et al., 2012). Consistently, viral RNAs, readily detected within pDCs, induce IFN response *via* the recognition by the endolysosome-localized TLR7 sensor (Decembre et al., 2014; Dreux et al., 2012; Takahashi et al., 2010). Of note, PI(4,5)P_2_, which clustered at contact (**Figure 4D-F**) is known to mark the plasma membrane and, together with cargo proteins, is instrumental for the initiation of clathrin-coated pit formation and consistently, PIP5KIα *via* PI(4,5)P2 synthesis control the rate and growth of clathrin-coated pit (Antonescu et al., 2011; He et al., 2017; Posor et al., 2015). We thus tested whether the components of the endocytosis machinery could polarize at contact and thereby facilitate the internalization of PAMP-carriers by pDCs. We observed co-clusters of clathrin and actin at the pDC/infected cell contact (**Figure 5A-C, 5G and S5A-D**). The 3D-reconstruction of consecutive Z-stack imaging suggested that the clathrin/actin co-clustering was present on the pDC side of the contact (**Figure 5C**). Similarly, Early Endosome Antigen 1 (EEA1), an endosomal marker, co-clustered with PI(4,5)P_2_ (**Figure 5D-F, 5H and S5E-F**) on the pDC side, as revealed by the 3D-reconstruction of contact sites (**Figure 5F**). The cell polarity of both these markers at contact required an intact actin network, as demonstrated by using inhibitor of actin polymerization (**Figure 5G-H**). Alike PI(4,5)P_2_/PIP5KIα co-clustering (**Figure 4G**), coclusters at contact of both clathrin/actin and EEA1/PI(4,5)P_2_ were less frequent as compared to the αL integrin (**Figure 2C**). Together our results suggest that components of the endocytosis machinery polarize at the contact between pDCs and infected cells, thus representing a functional remodeling to favor a localized endocytosis of PAMP-carriers leading to pDC activation *via* the endolysosomal TLR7.

**Figure 5.**
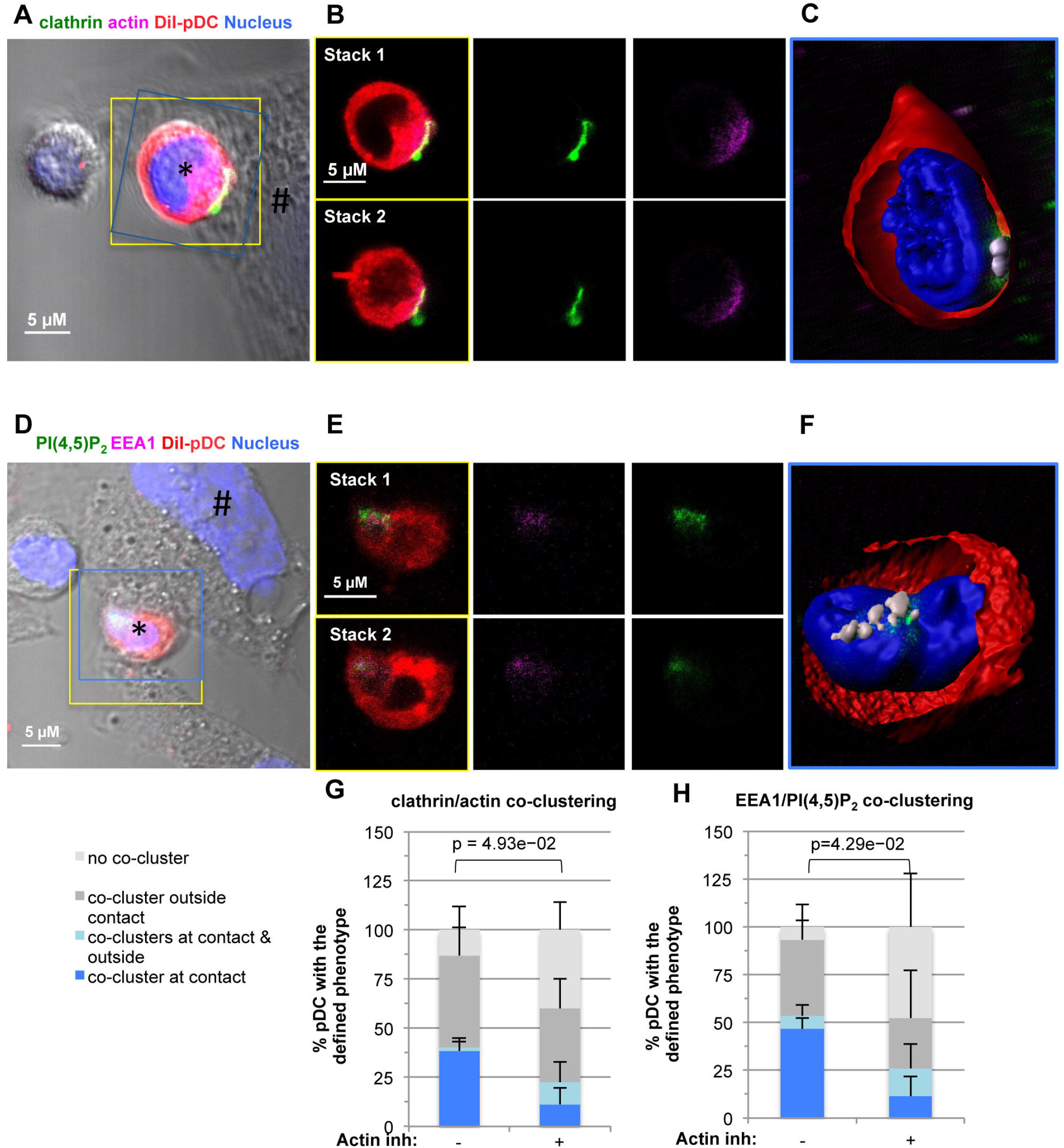
Polarization of endocytosis machinery components at the cell contact. **(A-C)** Confocal microscopy imaging of immunostained clathrin (green) and actin (purple) in 7-8 hour cocultures of pDCs (DiI stained, *) and HCV infected cells (#); nuclei (Hoechst). Representative detection of actin and clathrin projected on the phase contrast imaging (A), two successive confocal stacks with magnified yellow-boxed pDC (B) and 3D-reconstruction of consecutive Z-stacks of the blue-boxed view, with clathrin/actin colocalization in white (C). **(D-F)** Confocal microscopy imaging of immunostained EEA1 and PI(4,5)P_2_ co-clustering in 7-8 hour cocultures of pDCs and HCV replicating (SGR) cells, displayed as in panels A-C. **(G-H)** Quantification of clathrin/actin (G) and EEA1/PI(4,5)P_2_ co-clustering (H) in similar coculture treated or not with actin polymerization inhibitor (Latrunculin B, 1 mM), as in **Figure 4G**. Results of 4 (G; n>50 analyzed contacts per condition) and 3 (H; n>40 analyzed contacts per condition) independent experiments are expressed as the percentage of the total number of analyzed pDCs in contact as in **Figure 4G**; mean ± SD; p-value as indicated.

### pDCs establish sustained contact with virally infected cells as opposed to shorter contact with uninfected cells

The cell rearrangement at contact into a specialized territory enriched for regulators of actin network and endocytosis machinery was overall observed on the side of pDCs. These observations suggested the existence of a dynamic process within pDCs when in contact and sensing infected cells. We hypothesized that the sensing of PAMP-carriers released by infected cells by pDCs, and thereby the activated state of the pDCs could control the establishment of contact and/or the polarization of pDC cellular machineries at contact. We first tested whether sensing of the infected cells potentiated the establishment of cell contact. The quantitation of cell conjugates revealed that the frequency of pDCs in contact with infected cells was significantly higher as compared to uninfected cells (**Figure 6A-C**). Similar results were obtained for DENV and HCV infected cells, thus showing that the potentiation of contacts by sensing of virally infected cells is not virus-restricted. Additionally, increased frequency of contact was observed with HCV replicating cells, suggesting that the regulation is independent of the release of viral particle (**Figure 6C**).

**Figure 6.**
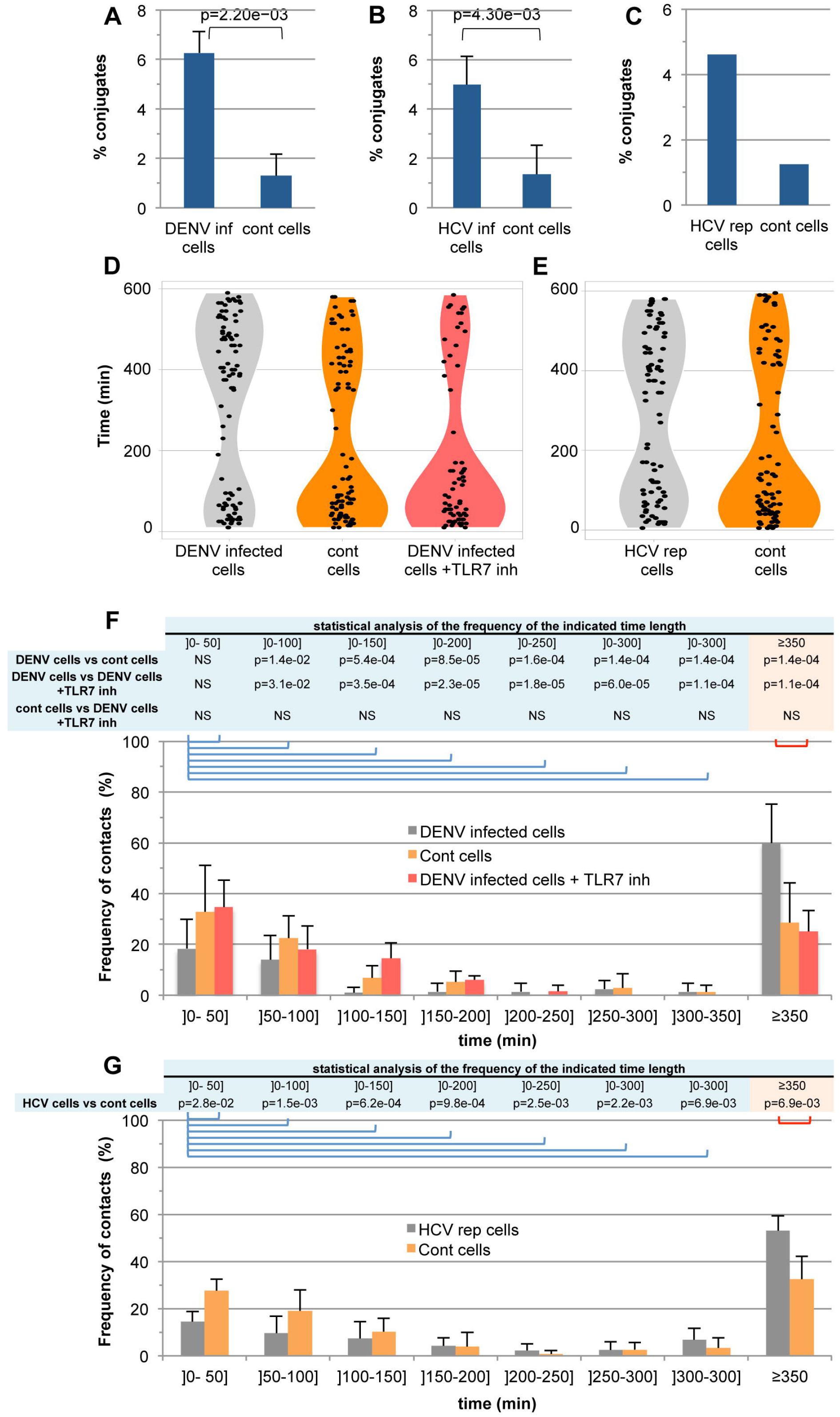
Establishment of sustained contact of pDCs with infected cells as opposed to shorter contact with uninfected cells. **(A-C)** Quantification of conjugates of pDCs with cells infected by DENV (A), HCV (B) and HCV replicating (rep) cells (C) as compared to uninfected cells by Image Stream X technology. Results of 6 (A, B) and 1 (C) independent experiments are expressed as percentage of conjugates; n=10 000 events per condition. **(D-G)** Quantification of the duration of cell contacts between pDCs and DENV infected cells **(D, F)** and HCV replicating (rep) cells **(E, G)**, determined by live-imaging with spinning-disc confocal microscopy. As indicated in panels **D** and **F**, cocultures were treated with TLR7 inhibitor (IRS661, 0.35 μM). Confocal Z-stacks of cocultures of DiI stained-pDC and GFP+ DENV infected cells or GFP+ HCV replicating cells are acquired every 5 minutes. The cell contact is defined as DiI signal (pDCs) apposed/merged with GFP signal (HCV/DENV cells), with visual inspection of randomly chosen fields, as shown on Figure S6. The results of 3-to-4 independent experiments are presented by violin plots with marks of durations of individual recorded contact in the indicated experimental conditions **(D-E)** and as percentages of contacts in the indicated time interval relative to the total number of examined pDC/infected cell contacts **(F-G)**. Data considered significant demonstrated adjusted p-value by False Discovery Rate (FDR) less than 0.05, as indicated.

The higher frequency of pDC contact with infected cells could reflect either contact with higher stability, which can better resist to the experimental procedure (*e.g*., collection of the cocultured cells) or an increased duration of the contact. To address these possibilities, we developed a live imaging-based approach using spinning-disk confocal microscopy analysis to quantify the duration of pDC contact with infected cells as compared to uninfected cells (**Figure S6A**). Distinct categories of contact duration were observed, including short-lived contacts (<100 min), intermediate duration contacts of low frequencies and sustained contact (>350 min) (**Figure 6D-G**). Importantly, the frequency of long contacts was higher when pDCs were cocultured with DENV infected cells, with reciprocal diminution of short contacts as compared to contact with uninfected cells (**Figure 6D** and F). In accordance, pDC mobility was reduced when cocultured with infected as compared to uninfected cells, as highlighted in tracking motion analysis (**Figure S6B**-E). Similar results were obtained in the context of pDC cocultured with HCV replicating cells (**Figure 6E** and 6G), suggesting this regulation is not virus-restricted and independent of the type of PAMP-carriers.

Our results demonstrated that pDCs established sustained contact with infected cells, thus implying that a molecular regulation of the contact by either sensing of the infected status of cells and/or the activating state of pDCs promotes their retention at the contact of infected cells. We thus tested the regulation of the pDC/infected cell contact by TLR7-induced signaling. To address this question, we used a specific TLR7 inhibitor (*i.e*., IRS661), which diminished the pDC IFNα production induced by DENV infected cells, as previously reported (Decembre et al., 2014). We found that the inhibition of TLR7 signaling diminished the frequency of the long-lived contacts of pDCs with infected cells with reciprocal higher number of short-lived contacts (**Figure 6D and F**). Inhibition of TLR7 signaling thus reverted the duration of pDC contacts to those observed for uninfected cells. Our results demonstrate that pDCs establish sustained contact with infected cells, via a feed-forward regulatory loop by TLR7-induced signaling.

### Recognition of viral infection favors polarization of cellular components at contacts via TLR7-induced signaling

Next, we tested whether the sustained contact observed with infected cells was associated with increased polarization of the identified cell adhesion molecules at the contact of pDCs with infected cells as compared to uninfected cells. The total surface expression of ICAM-1 did not markedly change when comparing infected *versus* uninfected cells (**Figure 1A and S1A**), neither did the level of α_L_ integrin on pDC cell surface when cocultured with either infected or uninfected cells (**Figure S7A**-B). Importantly, the co-accumulation at contact of α_L_integrin/ICAM-1 significantly augmented when pDCs were in contact with DENV or HCV infected cells as compared to uninfected cells (**Figure 7A-B**). The α_L_integrin/ICAM-1 interaction can induce mechanical transduction resulting in the polarization of actin network/regulators (DeMali et al., 2014). In accordance, we showed that the polarization of actin network regulators including PI(4,5)P_2_/PIP5KIα increased upon contact with HCV replicating cells (**Figure 7C**).

**Figure 7.**
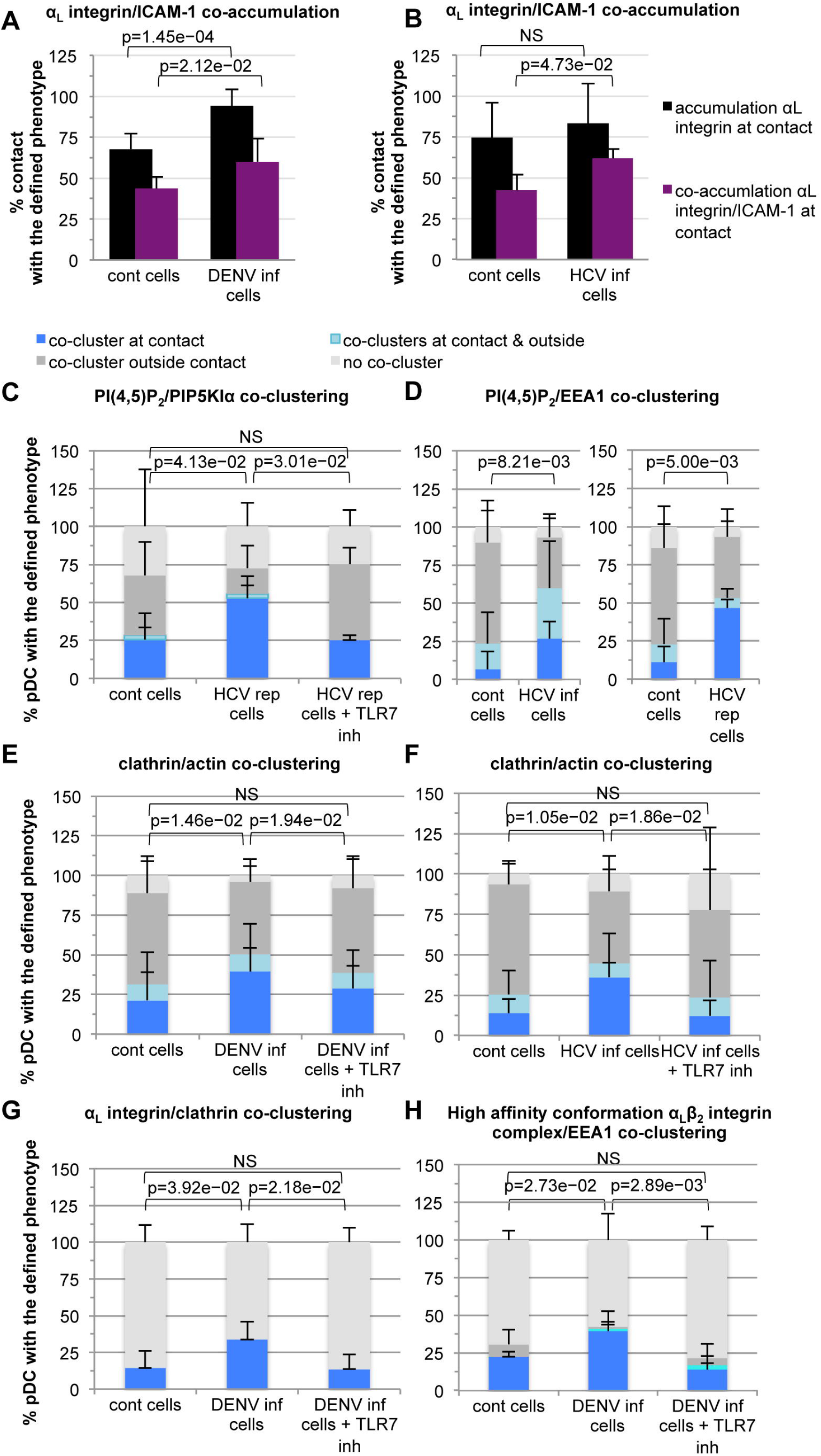
TLR7-mediated pDC activation potentiates cell polarity at the contact. **(A-B)** Quantification of the α_L_ integrin accumulation and α_L_ integrin/ICAM-1 co-accumulation at the contact between pDCs and DENV **(A)** or HCV (B) infected cells *versus* uninfected cells, analyzed as in **Figure 2I-K**. Results are expressed as the percentage of indicated phenotype relative to the total number of analyzed contacts; 4 (A; n>62 analyzed contacts per condition) and 3 (B; n>46 analyzed contacts per condition) independent experiments; mean ± SD; p-value as indicated. **(C-H)** Quantification of the co-clustering of PI(4,5)P_2_/PIP5KIα **(C)**, PI(4,5)P_2_/EEA1 **(D)**, clathrin/actin (E-F), α_L_ integrin/clathrin **(G)** and high-affinity conformation α_L_β_2_ integrin complex/EEA1 **(H)**, as shown in **Figure 4D-F, 5, S5, S7** in cocultures of pDCs with uninfected cells (cont cells), DENV infected, HCV infected and HCV rep cells, in the presence or not of TLR7 inhibitor (IRS661, 0.35 μM), as indicated. Results are expressed as the percentage of the indicated phenotype relative to the total number of analyzed pDCs in contact with infected cells; 3 (C-D), 7 **(E)**, 4 (F-H) independent experiments; mean ± SD; p-value as indicated.

We demonstrated that the inhibition of TLR7-induced signaling prevented the establishment of long-lived contact of pDCs with infected cells (**Figure 6D and F**). By analogy to inside-out signaling defined in the context of other cell-cell contact including neurological and immunological synapses (Hogg et al., 2011; Park and Goda, 2016), we thus hypothesized that TLR7-induced signaling might promote the cell polarity. We showed that the inhibition of TLR7 significantly reduced PI(4,5)P_2_/PIP5KIα co-clustering (**Figure 7C**), thus suggesting that TLR7-mediated signaling controls pDC polarity at contact. Similar regulation of cell polarity at contact was demonstrated for endocytosis components, including the coclustering of PI(4,5)P_2_/EEA1 and clathrin/actin (**Figure 7D-F**). The coclusters of α_L_ integrin and the high-affinity conformation of α_L_β_2_ integrin complex together with the endocytosis components (*i.e*., clathrin and EEA1, respectively) at the contact of pDCs with infected cells were also increased as compared to uninfected cells and subjected to regulation by TLR7-mediated pDC activation (**Figure 7G-H and S7C-D**). Similar results were obtained using DENV and HCV infected cells, highlighting that this feature of pDC functionality is neither virus-restricted nor related to the PAMP-carriers. Our results demonstrate that the sensing of infected cells *via* TLR7 sensor induces cell polarity of regulators of adhesion, actin network and endocytosis machinery at contact on the pDC side, thus acting as a positive feedback regulation for the efficient transfer of PAMP-carriers and response by pDCs.

## Discussion

pDCs are important mediators of IFN-I responses. We demonstrate that pDC response is mediated by the establishment of sustained contact with infected cells *via* cell adhesion molecules, α_L_β_2_ integrin complex and ICAM-1. This is associated with the polarization at the contact site and within pDCs, of the actin network and regulators as well as components of the endocytosis machinery, in accordance with the sensing by endolysosome-localized-TLR7. Furthermore, we provide compelling evidence that TLR7-induced signaling in pDCs regulates the rearrangement of a polarized platform at contact site and prolongs cell contacts, thus acting as a feed-forward regulation for the transfer of PAMP-carrier to the pDCs. Cell polarity is thus a previously unappreciated primary attribute of the pDCs to sense virally infected cells and respond by a robust antiviral response, which prevents viral spread. We propose to call this functional structuring the interferogenic synapse, by analogy to previously described synapses, while functionally driving a distinct cellular response.

### PAMP transfer *via* physical contact and cell polarity: the interferogenic synapse

Different types of cell-cell synapses have been defined including neurological, immunological and virological synapses. Although the required components, signals, cell types and functional outcomes vary, synapses have all in common to be cellular junctions specialized in the transfer of information from one cell to the other. Across all synapses, this information-sharing capacity is supported by adhesion complexes, thereby focusing signaling, secretion and endocytosis at the point of cell-cell contact (Dustin and Choudhuri, 2016; Hogg et al., 2011; Park and Goda, 2016). Likewise, we showed that adhesion molecules accumulate at the contact site of pDC/infected cell, and are required for pDC IFN response. Moreover, identical to the immunological synapse, the αLβ2 integrin complex is present on the signal recipient cells (pDCs *vs* T cells) and ICAM-1 on the signal delivering cells (infected cells *vs* antigen presenting cells) (Dustin and Choudhuri, 2016). Again alike all synapses, the actin network (and regulators) polarizes at contact and acts as a functional structuring platform. Analogous to the neurological synapse, the mechanism of pDC activation involved a secreted activating signal. Consistent with the transfer of PAMPs *via* vesicles and the endolysosomal localization of the sensor (*i.e*., TLR7), endocytosis components polarize at contact. Cell polarity of the vesicular trafficking is a hallmark of other synapses, primarily defined for secretory pathways, *e.g*., local secretion of synaptic vesicles or lytic granules at the neurological and immunological “lytic/NK” synapses, respectively. Additionally, the endocytosis machinery also polarizes at the level of the immunological synapse and is essential for T cell activation *via* the internalization and recycling of the T cell receptor (TCR) (Compeer et al., 2018; Das et al., 2004). Importantly, the feed-forward regulation by TLR7-induced signaling is reminiscent of the stabilization of the cell polarity and contact described for immunological and neurological synapses, referred to as inside-out signaling and induced by the response/activation in recipient cells (Dustin and Choudhuri, 2016; Hogg et al., 2011; Park and Goda, 2016). For example, T-cell receptor engagement at the immunological synapsed reinforces cell adhesion by the activation of integrin ectodomain (Dustin and Choudhuri, 2016; Hogg et al., 2011). Therefore, the interferogenic synapse presents hallmarks of previously described synapses, while being deployed to achieve a distinct cellular response, the IFN response, thus representing a newly described type of synapse.

### Scanning function of pDCs regulated by a TLR7-centered feed-forward signaling

We propose that the immune surveillance by pDCs involves a scanning function by physical cell-cell contacts (**Figure S8**). In this ‘scanning’ model, a pDC encounters a cell and initially establishes a transient contact. When the cell is not infected (*i.e*., in the absence of transfer of PAMPs), the non-activated pDC resumes its motion. By contrast, when a pDC encounters an infected cell, PAMP transfer during the initial contact activates TLR7 in the pDC. In turn, TLR7-mediated signaling potentiates the structural reorganization at the contact site (**Figure S8**). These molecular rearrangements include polarized endocytosis components, which can facilitate the subsequent transfer of PAMPs to pDCs and further strengthen TLR7 activation. TLR7-induced signaling primes the establishment of pDC/infected cell contact, which in turn further promote TLR7 activation, leading to a feed-forward regulatory loop culminating in sustained contacts as well as robust and localized IFN production. As such, the sensing mechanism by the pDCs can be seen as a “relay switch”: from a relatively small initial ‘PAMP current’, which lead to a ‘high amperage’ flow of PAMPs, by concentrating a functional platform for PAMP transfer at the contact sites, leading to a potent pDC-mediated antiviral response.

TLR7-induced signaling is expected to induce a bifurcated signaling: the transcriptional activators IRF7 and nuclear factor-κB (NF-kB), respectively leading to type I/III IFNs and other inflammatory cytokine response. One might hypothesize that the autocrine/paracrine action of IFNs (and/or subsequently induced-ISGs) and/or other inflammatory cytokine could mediate the potentiation of the cell contact and/or polarization of the pDCs. Alternatively, in accordance with previous publications (Eiseler et al., 2009; Olayioye et al., 2013; Park et al., 2009; Saitoh et al., 2017), the signaling downstream of TLR7, including the phosphorylation cascade *via* protein kinase D, a regulator of actin cytoskeleton and integrins might directly reinforce cell contact and pDC polarity.

### Local response to viral infection by pDCs

Systemic and massive productions of IFNs and other inflammatory cytokines are known to be detrimental to the host, as they correlate with systemic homeostatic dysfunction, including tissue damage and increased vascular permeability (*e.g*., via TNFα in the context of DENV (Costa et al., 2013)). Here we uncovered that the organization of functional territory in pDCs at the contact site with infected cells is required for the production of a large amounts of type I IFNs. The requirement of interferogenic synapse for a robust IFN response by pDCs thus implies that this antiviral response is confined to the proximity of infected cells. This is reminiscent of previous reports showing a pivotal *in vivo* antiviral regulation by early pDC response, despite undetectable or very transient/limited pDC-derived type I IFNs at systemic levels (Swiecki et al., 2010; Webster et al., 2018), thus positioning pDC type I IFN secretion as a response likely confined to the infected site. Therefore, the cell contact-dependent robust pDC response could have evolved in favor of the host fitness to locally respond at the infected sites, and thereby to thwart the otherwise harmful systemic IFN and inflammatory responses.

The requirement for cell contact with infected cells is increasingly recognized for pDC activation by a large spectrum of genetically divergent viruses (Webster et al., 2016). It is therefore likely that the interferogenic synapse that we describe here for pDC response to DENV, ZIKV and HCV *via* distinct PAMP-carriers, is also required for pDC IFN production triggered by other viruses. Additionally, pDCs can participate in the priming of both immunogenic and tolerogenic adaptive immunity, being also increasingly recognized as pivotal mediator in pathogenesis of autoimmune diseases and cancers (Swiecki and Colonna, 2015). It is conceivable that the interferogenic synapse is of importance for pDC-mediated sensing in these other contexts, such as *e.g*., autoimmune diseases displaying a prominent IFN signature induced by pDC TLR7/9 responses (Swiecki and Colonna, 2015).

## Acknowledgements

We thank A. D. Davidson (University of Bristol, Bristol, UK) for the molecular clone NGC, Alain Kohn (MRC-University of Glasgow Centre for Virus Research, UK) and Rafael Freitas de Oliveira Franca (FIOCRUZ, Recife, Brazil) for clinical isolate Brazil, PE243_KX197192), the European Virus Archive (EVA) for H/PF (H/PF/2013_KJ776791.1), C. Rice (Rockeffeler University, New York, NY) for the anti-HCV NS5A antibody (9E10), P. Despres (Pasteur Institut, Paris, France) for providing the anti-DENV E antibody (3H5), F.V. Chisari (Scripps Research Institute, La Jolla, CA) for the Huh7.5.1 and Huh7.5.1c2 cells and V. Lotteau (CIRI, Lyon, France) for Influenza A Virus. We are grateful to Y. Jaillais, H Paidassi, T. Walzer, S. Munoz-Gonzalez and C. Dong for critical reading of the manuscript, to S. This for assistance with graph preparation, and to our colleagues for encouragement and help. The PLATIM platform of SFR Biosciences Lyon-Gerland (UMS3444/US8) for confocal microscopy analysis and technical assistance. We are grateful to S. Dussurgey for assistance with Image-stream analysis. We acknowledge the contribution of the Etablissement Français du Sang Auvergne - Rhône-Alpes. This work was supported by grants from the ‘‘Agence Nationale pour la Recherche’’ (ANR-JCJC-EXAMIN), the ‘‘Agence Nationale pour la Recherche contre le SIDA et les Hépatites Virales’’ (ANRS-AO 2016-01) and the Fondation pour le recherche médicale (DBI20141231313 DREUX Bioinfo 2014). BW’s post-doc fellowship was sponsored by EMBO. SA’s PhD fellowship was sponsored by French ministry and Ligue contre le cancer.

## Author Contributions

Conceived and designed the experiments: SA, SC, WB, MD. Performed the experiments: SA, SC, ED, LS. Analyzed the data: SA, SC, ED, LS, OA, BW, MD. Wrote the paper: SA, SC, MD.

## STAR Methods

### Biological materials

Huh7.5.1, Huh-7.5.1c2 cells and Huh-7.5.1c2 cells that harbor the JFH-1 subgenomic replicon (HCV rep cells) (Kato et al., 2003) were maintained in Dulbecco’s modified Eagle medium (DMEM) (Life Technologies) supplemented with 10% FBS, 100 units (U)/ml penicillin, 100 mg/ml streptomycin, 2 mM L-glutamine and non-essential amino acids (Life Technologies) at 37°C/5% CO_2_ (Decembre et al., 2014; Dreux et al., 2012). The pDCs were isolated from blood or cytapheresis units from healthy adult human volunteers which was obtained according to procedures approved by the “Etablissement Français du sang” (EFS) Committee. PBMCs were isolated using Ficoll-Hypaque density centrifugation and pDCs were positively selected from PBMCs using BDCA-4-magnetic beads (MACS Miltenyi Biotec), as previously described (Decembre et al., 2014; Dreux et al., 2012). The typical yields of PBMCs and pDCs were 800×10^6^ and 2×10^6^ cells, respectively, with a typical purity of >95% pDCs. Isolated pDCs were maintained in RPMI 1640 medium (Life Technologies) supplemented with 10% FBS, 10 mM HEPES, 100 units/ml penicillin, 100 mg/ml streptomycin, 2 mM L-glutamine, non-essential amino acids and 1 mM sodium pyruvate at 37°C/5% CO_2_. The down regulation of ICAM-1 in huh7.5.1 cells was achieved using lentivirus-based vector expressing shRNA against ICAM-1 (TRCN0000372477, target sequence: GCCCGAGCTCAAGTGTCTAAA, Sigma-Aldrich), produced as we previously described (Dreux et al., 2012). Cells were transduced 4 days prior to infection by DENV for 48 hours and followed by coculture with pDCs.

### Preparation of viral stocks and infections

Viral stocks of HCV JFH1 strain (Zhong et al., 2005) and the prototypic DENV-2 strain New Guinea C (NGC) (AF038403) (Kroschewski et al., 2008) were produced as previously described (Decembre et al., 2014; Dreux et al., 2012). The clinical isolate of epidemic strains of ZIKV including from Brazil (PE243_KX197192) (Donald et al., 2016) and H/PF from French Polynesia (H/PF/2013_KJ776791.1) obtained from the European Virus Archive (EVA) were amplified using Vero cells. Huh7.5.1 cells were infected by DENV or ZIKV (MOI of 1) and Huh-7.5.1c2 cells by the cell-culture adapted HCV JFH-1 virus (MOI of 1) (Decembre et al., 2014; Dreux et al., 2012) 48 hours prior to co-culture with isolated pDCs.

Viral stocks of Influenza A Virus (FluAV, H1N1/New Caledonia/2006) were produced as previously described (de Chassey et al., 2013) and kindly provided by Dr V. Lotteau (CIRI, Lyon, France).

### Reagents

The antibodies used for immunostaining were mouse anti-DENV E glycoprotein (3H5) kindly provided by P. Despres (Pasteur Institut, Paris, France); mouse anti-HCV NS5A (clone 9E10) kindly provided by C. Rice (Rockefeller Université, New York, US); mouse anti-dsRNA (clone J2; Scicons); mouse anti-integrin α_L_ subunit (clone 38; LSBio); mouse anti-integrin β_2_ I-like domain, *i.e* high-affinity conformation - epitope m24 (Biolegend); mouse anti-ICAM-1 (clone LB-2; BD Pharmingen); mouse anti-E-Cadherin (SHE78-7; Life Technologies); mouse anti-L-selectin/CD62L (unconjuguated clone DREG55; Invitrogen and Clone 145/15; Vioblue-conjugated; Miltenyi); mouse anti-actin (clone AC74; Sigma-Aldrich); rabbit anti-protein disulfide isomerase (PDI) (anti-P4HB, HPA018884, Sigma-Aldrich); mouse anti-PI(4,5)P_2_ (ab2335, Abcam); goat anti-PIP5KIα (reference C-17/sc-11774, Santa Cruz); rabbit anti-clathrin (ab21679, Abcam); rabbit anti-EEA1 (ALX-210-239, Enzo Life Sciences); pDC specific markers: mouse PE or APC-conjugated anti-CD123 (clone AC145, Miltenyi), mouse APC-conjugated anti-BDCA-2 (AC144; Miltenyi) and Alexa-conjugated secondary antibodies (Life Technologies).

The inhibitors used were Latrunculin B (Sigma-Aldrich); Arp2/3 complex inhibitor I (CK-666, Merck Millipore); CDC42 inhibitor (ML141, Sigma-Aldrich); TLR7 inhibitor IRS661 (5’-TGCTTGCAAGCTTGCAAGCA-3’) synthesized on a phosphorothioate backbone (Decembre et al., 2014).

The other reagents were TLR7 agonist (IMQ; Imiquimod, Invivogen); Ficoll-Hypaque (GE Healthcare Life Sciences); Fc Blocking solution (MACS Miltenyi Biotec); IFNα ELISA kit (PBL Interferon Source); 96-Well Optical-Bottom Plates and Nunc UpCell 96F Microwell Plate (Thermo Fisher Scientific); 96-well format transwell chambers separated by a 0.4 mm membrane (Corning); Vibrant cell-labeling solution (CM-DiI, Life Technologies); Celltrace Violet cell Proliferation kit (Life Technologies); Hoescht and Fc receptor blocking reagent (MACS Miltenyi Biotec); cDNA synthesis and qPCR kits (Life Technologies); poly-L-lysin (P6282, Sigma-Aldrich).

### Immunostaining and FACS analysis

Surface immunostainings were performed on freshly isolated pDCs, DENV infected and uninfected Huh-7.5.1 cells and cocultured cells (Figure S6) that were harvested and resuspended using 0.48 mM EDTA-PBS solution. Surface adhesion molecules were detected by a 30 minute incubation at 4°C with 5 μg/mL of mouse anti-α_L_ integrin (clone 38); mouse anti-ICAM-1 (Clone LB-2); mouse anti-E-cadherin (clone SHE78-7); Vioblue-conjugated mouse anti-L-selectin/CD62L (clone 145/15) diluted in staining buffer (PBS - 1% FBS), followed by PBS washes. For α_L_ integrin, ICAM-1 and E-cadherin detection, cells were incubated for 20 minutes with APC-conjugated-anti-mouse antibody diluted in staining buffer. Flow cytometric analysis was performed using a Digital LSR II, and the data were analyzed with Flow Jo software (Tree Star).

### Co-culture experiments

Unless otherwise indicated, 10^4^ pDCs were co-cultured with 2×10^4^ infected cells or uninfected parental cells in a 200 μl final volume in 96-well round-bottom plates incubated at 37 °C/5% CO_2_. Eighteen to twenty hours later, cell-culture supernatants were collected and the levels of IFNα was measured using a commercially available ELISA kit specific for IFNα (PBL Interferon Source) following the manufacturer’s instructions. To optimize the pDC response to HCV replicating (SGR) cells, 2×10^4^ pDCs were co-cultured with 10^5^ cells for 22-to-24 hours.

### Viral spread assay

5×10^4^ GFP-expressing uninfected cells and 5×10^4^ GFP-negative cells, that were infected by DENV or ZIKV, as indicated, for 48 hours prior to coculture, were cocultured with 2×10^4^ pDCs, 48 hour-incubation. When indicated, the cocultures were treated by anti-α_L_ integrin at 10 μg/mL. The levels of the viral spread from infected cells (GFP^-^) to uninfected cells (GFP^+^) during coculture were determined by FACS analysis of the frequency of infected cells in the GFP^+^ cell population. Similar detection in GFP^-^ populations demonstrated the inhibition of viral replication during coculture. The infected cells were detected by immunostaining of dsRNA, a marker of viral replication. Briefly, after 48 hours of coculture, cells were fixed with 4%PFA, followed by PBS washes and a permeabilization step with 90% methanol/10% PBS for 1 hour-incubation at 20°C. The mouse anti-dsRNA (clone J2; Scicons) was diluted staining buffer (PBS - 1% FBS) and added to the cells for an hour incubation at 4°C. After PBS wash, cells were incubated with anti-mouse IgG2a conjugated with Alexa 647-fluorochrome and diluted staining buffer with 30 minute incubation at 4°C. Flow cytometric analysis was performed using a Digital LSR II, and the data were analyzed with Flow Jo software (Tree Star).

### Imaging combined with flow cytometry analysis of pDC/infected cell conjugates using Image Stream X technology

Huh 7.5.1c2 cells were transduced with lentiviral based vector pseudotyped with VSV glycoprotein to stably express GFP, as previously described (Decembre et al., 2014; Dreux et al., 2009). Forty-eight hours prior co-culture with pDCs, GFP-expressing Huh 7.5.1c2 cells were infected by HCV or DENV as above-described. After immuno-isolation, pDCs were stained by using Celltrace Violet Cell Proliferation kit (Life Technologies) by 20 minute-incubation at 37°C in the dark. Labeled pDCs were then spinned down and resuspended into pDC coculture medium. 10^5^ GFP-expressing HCV/DENV infected or uninfected cells were co-cultured with 4 x 10^4^ pDCs in low-adherence micro-plate designed for cell harvesting by temperature reduction (Nunc UpCell 96F Microwell Plate from Thermo Scientific) for 4-5 hours at 37°C in presence or not of inhibitor, as indicated. The co-cultured cells were detached and harvested by 20 minute-incubation at room temperature. After 2% PFA fixation, cells are washed twice with staining buffer (PBS 2% FBS). Co-cultured cells were analyzed by Image Stream X technology (Amnis) at magnification x40 using IDEAS software. The cell population defined as pDC/HCV-infected or uninfected cell conjugates comprises conjugates of at least one Celltrace Violet (CTV) positive cell and at least one GFP positive cell among the total of CTV positive cells, GFP positive cells and conjugates. The cell populations were sorted by using masks (IDEAS software) to eliminate cells not in focus and/or with saturating fluorescent signal, conjugate population was further selected by with Area dilate IDEAS mask (*i.e*., double fluorescent signal without overlay) and then sorted based of cell size of the positive cells (*i.e*., fluorescent signal area). Post-cell sorting, the accuracy of the gated cell populations in regards to the defined criteria was controlled by a visual inspection of the individual pictures in the gated cells population (**Figure S1B-C**). When indicated, after 4% PFA fixation, cells were incubation with Fc receptor blocking reagent (MACS Miltenyi Biotec) for 10 minutes at 4°C, followed by a 40 minute incubation at 4°C with 5 μg/mL of APC-conjugated mouse anti-CD123 (pDC marker), diluted in staining buffer, followed by washes with staining buffer and analyzed by Image Stream X technology.

### Immunostaining for confocal analysis

After immuno-isolation, 4 x 10^4^ pDCs were stained by using either 0.5 μM Vibrant cell-labeling solution (CM-DiI, Life Technologies) by successive incubations for 10 and 15 minutes at 37°C and 4°C, respectively or Celltrace Violet cell Proliferation kit (Life Technologies) as above-described. Labeled pDCs were washed twice with PBS and then co-cultured with 2 x 10^4^ DENV infected Huh7.5.1 cells or HCV infected Huh7.5.1c2 cells at 37°C for 4-5 hours, in a 96-Well Optical-Bottom Plate precoated with 8 μg/mL poly-L-lysin for 30 minutes incubation at 37°C. A longer incubation time (*i.e*., 7-8 hours) was preferred for analysis of clathrin/actin and EEA1/PI(4,5)P_2_ in HCV cells, as a pilot experiment suggested that the polarization of endocytic components is slightly higher at this time point. After 4% PFA fixation and three PBS washes, immunostainings were performed without additional permeabilization step, as previously (Decembre et al., 2014; Meertens et al., 2012). After blocking step (PBS 3% BSA), primary antibody antibodies were diluted in 3% BSA-PBS and added to the cell for one hour incubation at room temperature. Primary antibodies used in this study include mouse anti-actin (20 μg/mL); rabbit anti-PDI (2 μg/mL); mouse anti-PI(4,5)P_2_ (20 μg/mL); goat anti-PIPK1α (4 μg/mL); rabbit anti-clathrin (2 μg/mL); rabbit anti-EEA1 (2 μg/mL); mouse anti-ICAM-1 (10 μg/mL); mouse anti-α_L_ integrin subunit (clone 38 LSBio at 5 μg/mL); mouse anti-integrin β_2_ I-like domain, *i.e*., open/high-affinity conformation of α_L_β_2_ integrin complex (clone m24 at 5 μg/mL); APC-conjugated mouse anti-CD123 (clone AC145 at 5 μg/mL). After three washes with PBS, cells were incubated with the appropriate anti-mouse, rabbit or goat antibody [conjugated with Alexa 488-, 555- or 647-fluorochrome] or Alexa-conjugated antibody against specific mouse isotypes *[i.e*., anti-IgG1 for detection of m24 epitope of a_L_β_2_ integrin complex and anti-IgG2a for detection of the integrin α_L_ subunit] at 2 μg/mL in 3% BSA-PBS and add to the cells along with Hoechst diluted at 1:1000 (Molecular Probes) for 1 hour incubation at room temperature. After three gentle washes with PBS, cells were observed with a Zeiss LSM 710 laser scanning confocal microscope. The quantification of the phenotypes defined to as clusters or co-clusters at the contact, outside contact or no cluster/co-clusters were performed using Image J software package (http://rsb.info.nih.gov/ij); Results were validated in a “double-blind” set-up. IMARIS (Bitplane Inc.) software package with coloc tools was used for the 3D-reconstructions of the consecutive Z-stacks.

### Live imaging of coculture with spinning-disc confocal microscopy analysis

Huh 7.5.1c2 cells were transduced with lentiviral based vector to stably express GFP and infected with HCV or DENV 48 hours before co-culture, as above described. GFP-expressing DENV infected cells or HCV replicating cells and corresponding uninfected control (cont) cells were seeded (2 x 10^4^ cells per well) in a 96-Well Optical-Bottom Plate pre-coated with poly-L-lysin (30 minutes incubation at 37°C with 8 μg/mL poly-L-lysin). Isolated pDCs were stained with Vibrant cell-labeling solution (CM-DiI, Life Technologies) as above described. When pDCs were added to the seeded infected (or uninfected) cells, the cocultures were imaged every 5 minutes for 10 hours at magnification ×10 with a BSL3-based spinning-disc confocal microscope (AxioObserver Z1, Zeiss). The analyses of pDC motion (as illustrated in Figure S6) were performed using projection of Z-stacks with maximal intensity (*i.e*., about 3-to-5 selected Z-stacks per fields out of 15 Z-stacks in total). The quantification of the duration of contacts between pDCs and infected cells versus uninfected cells were performed using Image J software package (http://rsb.info.nih.gov/ij). The representation with violin plots were obtained using R software environment for statistical computing and graphics, version 3.3.2. The calculations of position/motion for the motion graphs were performed using Trackmate plug-in of Image J software.

### Analysis of intracellular and extracellular RNA levels

RNAs were isolated from cells or supernatants harvested in guanidinium thiocyanate citrate buffer (GTC) by phenol/chloroform extraction procedure as previously described (Dreux et al., 2012). The efficiency of RNA extraction and reverse transcription-real-time quantitative PCR (RT-qPCR) was controlled by the addition of carrier RNAs encoding Xef1α (xenopus transcription factor 1α) *in vitro* transcripts in supernatants diluted in GTC buffer. DENV and HCV RNA, Xef1α and intracellular glyceraldehyde-3-phosphate dehydrogenase (GAPDH) mRNA levels were determined by RT-qPCR using cDNA synthesis and qPCR kits (Life Technologies) and analyzed using StepOnePlus Real-Time PCR system (Life Technologies), using previously described primers (Decembre et al., 2014; Dreux et al., 2009). Extracellular and intracellular DENV and HCV RNA levels were normalized for Xef1α and GAPDH RNA levels, respectively.

### Analysis of extracellular infectivity

Infectivity titers in supernatants were determined by end-point dilution using Huh 7.5.1 cells. Foci forming unit (ffu) were detected 72 hours after infection by anti-HCV NS5A and anti-DENV E glycoprotein specific immunofluorescence for HCV and DENV, respectively, as previously described (Decembre et al., 2014; Dreux et al., 2009). Briefly, Huh-7.5.1 cells were fixed with 4% PFA and permeabilized by incubation for 7 minutes in PBS containing 0.1% Triton. Cells were then blocked in PBS containing 3% BSA for 15 minutes and incubated for 1 hour with mouse anti-HCV NS5A (clone 9E10) diluted at 1:500 or mouse anti-DENV E glycoprotein (clone 3H5) hybridoma supernatant diluted at 1:200 in PBS containing 1% BSA. After 3 washes with PBS, cells were incubated 1 hour with secondary Alexa 555-conjugated anti-mouse antibody (2 μg/mL) and Hoechst dye in PBS containing 1% BSA. Percentage of NS5A-positive and E-positive cells cells was determined using Zeiss Axiovert 135 or Olympus CKX53 microscopes.

### Statistical analysis

Statistical analysis was performed using R software environment for statistical computing and graphics, version 3.3.2. For levels of IFNα productions, viral RNA levels, viral titers and cell surface expression, the values were considered relative to untreated references for each independent experiment and analysis using an one-way ANOVA on ranks (Kruskal-Wallis test). When the test was significant, we used the Nemenyi *post hoc* test for multiple comparisons of mean rank sums to determine which contrast(s) between individual experimental condition pairs was significant. For the statistical analysis of polarization of structure/component at contact (assessed by confocal microscopy analysis), the distribution of the analyzed pDCs in contact with infected cells that presents co-clusters at the contact (without or with additional cluster outside contact) *versus* no co-cluster at contact (without or with cluster outside contact) were compared using Fisher’s exact test for count data. Similar results were obtained when using the test for equality of proportions. The results of imaging flow cytometry (using Image Stream X technology) were analyzed using the Wilcoxon Unpaired test. For statistical analysis of the duration of contact (assessed by live-imaging using spinning-disk confocal microscopy analysis), a test for pairwise equality of proportions was applied to determine whether the duration of contact depends on the experimental conditions by comparing the frequency of the indicated time lengths. Data considered significant demonstrated adjusted p-value by False Discovery Rate (FDR) less than 0.05.

## Supplementary Figure Legends

**Figure S1, related to **Figure 1**. Surface expression of adhesion molecules and impact of their inhibition on pDC IFNα production and the frequency of pDC/infected cell conjugates. (A)** FACS analysis of adhesion molecule expression at the cell surface of pDCs, DENV infected and uninfected (cont) Huh7.5.1 cells. Representative histograms with unstained cells or stained only with secondary antibody as reference for immunostained L-selectin or E-cadherin/α_L_ integrin /ICAM-1, respectively; representative of 3-4 independent experiments. **(B)** Quantification of IFNα in supernatants of pDCs cocultured with DENV infected cells, treated or not with blocking antibodies against α_L_ integrin or ICAM-1, at the indicated concentration. The intracellular and extracellular viral RNA levels and infectious virus production by infected cells (in absence of pDCs) treated with blocking antibodies were analyzed. Results and statistical significances are presented as in **Figure 1B**; mean ± SD; n=4-7 independent experiments. **(C-D)** DENV infected Huh7.5.1 cells, which stably express GFP marker (green), were cocultured with pDCs for 4-5 hours. pDCs are detected by staining with CellTrace Violet Cell Proliferation solution prior to coculture (purple). **(C)** Representative pictures of the cell populations gated as conjugates between pDCs and GFP expressing DENV infected cells (right panel) as well as single cell populations (right panels). Panels, as displayed from the left to the right: Bright field; GFP field; Celltrace-stained pDC field and Merge. Of note, we did not observed any cell double positive for GFP and Celltracer Violet Cell Proliferation for 5071 pDC-infected cell conjugates; 7 independent experiments, with visual validation, thus assessing the absence of cytosolic mixing between the infected cells and pDCs. **D)** Visual examination of the individual images from the population gated as conjugates. Results are expressed percentage of conjugates in population gated as conjugates; analysis 6 independent experiments with indicated cell numbers for each type of pDC staining.

**Figure S2, related to Figure 2. Accumulation of α_L_β_2_ integrin is observed at pDC/infected cell contact, but not in absence of contact. (A-B)** Confocal microscopy imaging of immunostained α_L_ integrin (green) and CD123 (purple) in co-cultures of pDCs (stained by the lipophilic dye DiI prior coculture; red; star mark *) and DENV infected cells (#) for 4-5 hour incubation, nuclei stained with Hoechst (blue), as in **Figure 2A-B**, with detection of αL integrin and CD123 projected on the phase contrast imaging (A) and magnification of two consecutive Z-stacks of yellow-boxed pDC (B). **(C-D)** Confocal microscopy imaging of immunostained open/high-affinity conformation of α_L_β_2_ integrin complex (m24 epitope; green) in co-cultures of pDCs (DiI stained; red; *) and DENV infected cells (#), 4-5 hour-incubation, nuclei (Hoechst, blue), displayed as in **Figure 2D-E. (E-F)** Confocal microscopy imaging of immunostained αL integrin (green) in absence of contact of pDCs (*) with infected cells, displayed as in panel **A-B.**

**Figure S3, related to **Figure 3**. Inhibition of viral propagation by pDC response to contact with ZIKV-infected cells**. FACS analysis of viral transmission from GFP^-^ ZIKV infected cells to GFP+ uninfected cells when cocultured with pDCs (48 hours incubation) in presence of anti-αL integrin (10 μg/mL), as indicated. ZIKV infected cells are detected by immunostaining of the dsRNA viral marker. Results presented as dot plots are representative of 3 independent experiments. The frequencies of the cell populations are indicated as percentages on the dot plot.

**Figure S4, related to Figure 4. Actin network polarizes at contact and is required for pDC response. (A-C)** Confocal microscopy imaging of immunostained actin (purple) and the ER marker PDI (green) in co-cultures of pDCs (DiI stained, *) and HCV infected cells (#), 4-5 hour incubation; nuclei (Hoechst, blue). Detection of actin and PDI projected on the phase contrast imaging (A), two successive confocal stacks with magnified yellow-boxed pDC (B) and quantification of the frequency of pDCs with clustering of actin and PDI at contact (C). Results of 3 independent experiments are expressed as the percentage relative to the total number of analyzed pDCs in contact with HCV infected cells, mean ± SD; n=31 analyzed contacts par condition; p-value as indicated. **(D-E)** Quantification of IFNα in supernatants of pDCs cocultured with DENV infected (D) or HCV replicating (rep) cells (E) that were treated or not with CDC42 inhibitor (ML141, at the indicated concentrations). The intracellular and extracellular viral RNA levels were analyzed for infected cells similarly treated with inhibitors (in absence of pDCs). Mean ± SD; n=3-6 independent experiments; p-value as indicated; non-detectable (arrays). **(F)** Quantification of IFNα in supernatants of pDCs cocultured with HCV infected or HCV replicating (rep) cells that were treated or not with inhibitor of actin polymerization (Latrunculin B, 1 mM), 2×10^4^ pDCs and 10^5^ HCV cells for 22-to-24 hours. The intracellular and extracellular viral RNA levels of HCV infected/replicating cells (in absence of pDCs) upon treatment conditions (*i.e*., incubation time and concentration) exactly as for the analysis of IFNα production were analyzed. Results are expressed as percentages relative to untreated cells; mean ± SD; n=4 independent experiments; non detectable (arrays). **(G)** Quantification of pDC/infected cell conjugates by Image Stream X technology. pDCs cocultured with GFP expressing HCV infected or replicating (rep) cells for 4-5 hours in presence or not actin polymerization inhibitor as in panel F. Cocultures were stained using the pDC specific marker CD123 prior to analysis by Image Stream X technology. Results of 5 (HCV inf cells) and 1 (HCV rep cells) independent experiments expressed as percentage of conjugate; mean ± SD; n=10 000 events per condition; p-value as indicated.

**Figure S5, related to Figure 5 and 7. Co-clustering of regulators of actin network and endocytosis machinery at the contact. (A-D)** Confocal microscopy imaging of immunostained actin (purple) and clathrin (green) in cocultures of pDCs (DiI-stained, *) and HCV infected cells (A-B) or DENV infected cells (C-D); nuclei (Hoechst). Representative detection of actin and clathrin projected on the phase contrast imaging (A and C) and two successive confocal stacks with magnified yellow-boxed pDCs (B and D). **(E-F)** Confocal microscopy imaging of immunostained EEA1 and PI(4,5)P_2_ in cocultures of pDCs (DiI-stained, *) and HCV infected cells; nuclei (Hoechst). Representative detection of EEA1 and PI(4,5)P2 projected on the phase contrast imaging (E) and two successive confocal stacks with magnified yellow-boxed pDC (F).

**Figure S6, related to Figure 6. Live-imaging analysis of the contacts between pDCs and infected cells versus uninfected cells. (A)** Representative live imaging of DiI stained-pDC (red) cocultured with GFP expressing DENV infected cells (green) analyzed using spinning disc confocal microscopy. Results are presented as a series as 5 min interval imaging, *i.e*.,confocal Z-stacks projected on phase contrast imaging; motion tracking of pDCs as indicated in yellow; red frames corresponding of time of the pDC/infected cells contact are highlighted in red. **(B)** Representative confocal Z-stacks imaging of cocultures of DiI stained-pDC (red) and GFP expressing DENV infected cells (green) projected on phase contrast imaging, with indication (numbering) of the pDCs analyzed in motion tracking in (C). **(C)** Motion tracking graph of 10 randomly selected pDCs cocultured with infected cells. Time-lapse images were processed using Trackmate plug-in of Image J software, and position/motion of each individual tracked pDC reported on the graph. **(D)** Representative imaging of cocultures of DiI stained-pDC GFP expressing uninfected (cont) cells, presented as in B. **(E)** Motion tracking graph of 10 randomly selected pDCs cocultured with uninfected cells, presented as in **C.** All live-imaging of cocultures with infected and uninfected cells were performed in parallel.

**Figure S7, related to Figure 7. Surface expression of α_L_ integrin on cocultured pDCs and imaging of co-clustering of the adhesion and vesicular trafficking components at the contact. (A-B)** FACS analysis of immunostained αL integrin at the cell surface of pDCs cocultured with DENV infected Huh7.5.1 cells *versus* uninfected (cont) cells for 4-5 hours. Results are presented with representative histogram (A) and quantification of the surface expression (B), expressed as fold-increases of geomean relative to cells stained only with secondary antibody; fold-increase geomean ± SD; n=3 independent experiments. **(C)** Representative confocal microscopy imaging of immunostained αL integrin (green) and clathrin (purple) in cocultures of pDCs (CTV-stained;*) and DENV infected cells; nuclei (Hoechst); detection of α_L_ integrin/clathrin projected on the phase contrast imaging (left panels); two successive confocal stacks with magnified yellow-boxed pDC (middle panels) and 3D-reconstruction of consecutive Z-stacks of the blue-boxed view (right panel). **(D)** Representative confocal microscopy imaging of immunostained high-affinity conformation of α_L_β_2_ integrin complex (*i.e*., m24 epitope) and the early endosome EEA1 marker in cocultures of pDCs (DiI-stained, *) and DENV infected cells (#); nuclei (Hoechst); detection of high-affinity conformation α_L_β_2_ integrin complex and EEA1 projected on the phase contrast imaging (left panels), two successive confocal stacks with magnified yellow-boxed pDC (middle panels) and 3D-reconstruction of consecutive Z-stacks of blue-boxed pDCs (right panel).

**Figure S8. Working model of the immune surveillance by pDCs by a scanning function thought cell contacts.** The pDC could scan the cells by non-stabilized cell contact. If the cell is not infected (in green), the non-activated pDC will resume its motion. In contrast, when the pDC encounters an infected cell (in red), PAMP transfer during the initial contact activates TLR7 sensor, and subsequent structural reorganization at the contact site within pDC. As shown in the zoomed inset of the schematic representation of the cell contact, the molecular rearrangement at the contact site includes the polarization of cell adhesion complex (*i.e*., ICAM-1/α_L_β_2_ integrin) and regulators of actin network (*i.e*., Arp2/3, CDC42, PI4,5P_2_, PIP5K1α) and endocytosis machinery. The latter can facilitate the subsequent transfer of PAMPs to pDCs and further strengthen TLR7 activation, leading to a feed-forward regulation culminating in sustained contacts and a robust IFN production by the pDCs.

